# Reconstruction of post-synaptic potentials by reverse modeling of local field potentials

**DOI:** 10.1101/346148

**Authors:** Maxime Yochum, Julien Modolo, Pascal Benquet, Fabrice Wendling

## Abstract

Among electrophysiological signals, Local Field Potentials (LFPs) are extensively used to study brain activity, either *in vivo* or *in vitro*. LFPs are recorded with extracellular electrodes implanted in brain tissue. They reflect intermingled excitatory and inhibitory processes in neuronal assemblies. In cortical structures, LFPs mainly originate from the summation of post-synaptic potentials (PSPs), either excitatory (ePSPs) and inhibitory (iPSPs) generated at the level of pyramidal cells. The challenging issue, addressed in this paper, is to estimate, from a single extracellularly-recorded signal, both ePSP and iPSP components of the LFP. The proposed method is based on a model-based reverse engineering approach in which the measured LFP is fed into a physiologically-grounded neural mass model (mesoscopic level) in order to estimate the synaptic activity of a sub-population of pyramidal cells interacting with local GABAergic interneurons. The method was first validated using simulated LFPs for which excitatory and inhibitory components are known *a priori* and can thus serve as a ground truth. It was then evaluated on *in vivo* data (PTZ-induced seizures, rat; PTZ-induced excitability increase, mouse; epileptiform discharges, mouse) and on *in clinico* data (human seizures recorded with depth-EEG electrodes). Under these various conditions, results showed that the proposed reverse engineering method provides a reliable estimation of the average excitatory and inhibitory post-synaptic potentials at the origin of the measured LFPs. They also indicated that the method allows for monitoring of the excitation/inhibition ratio. The method has potential for multiple applications in neuroscience, typically when a time tracking of local excitability changes is required.

## 1. Introduction

Local field potentials (LFPs) refer to extracellularly-recorded electrophysiological signals generated by neuron assemblies (Buzsáki 2004). LFPs are extensively used to monitor and analyze brain function in brain research as well as in neurological clinical studies, Since LFP recordings require minimal equipment and provide critical information on local neuronal activity, they are used in a wide spectrum of applications, from basic neuroscience to clinics (Destexhe & Bedard 2013) (e.g., using the LFP as a control signal to trigger stimulation in closed-loop (Priori et al. 2013)).

Physiologically, LFP signals reflect ongoing excitation- and inhibition-related processes in recorded networks (Buzsáki et al. 2012). Indeed, it is known that the major contribution to LFPs arises from post-synaptic potentials (PSPs) generated at the level of pyramidal cells within spatially extended neuronal assemblies. As described by bioelectromagnetic models (Malmivuo et al. 1995, Ramachandran 2002) and experimental studies (Buzsáki et al. 2012), the summation of PSPs, either excitatory (ePSPs) or inhibitory (iPSPs) generated at the level of pyramidal cells located in the cerebral cortex is the major contribution to LFPs. This is essentially explained by two factors: (1) synaptic activation leads to the formation of a sink and a source at the level of neurons, which can then be viewed as elementary current dipoles (Buzsáki et al. 2012), and (2) when neurons are geometrically aligned (Buzsáki et al. 2012, Harris et al. 2003) (like pyramidal cells organized “in palissade” in cortical structures), then dipole contributions tend to sum up instead of cancelling out (Buzsáki et al. 2012). In addition, due to the frequency-filtering properties of cerebral tissue, slow post-synaptic currents are more likely to be recorded further away from the electrode contacts, as opposed to fast transmembrane ionic currents (involved in action potentials) which attenuate much faster with space (Bédard et al. 2004). Overall, LFPs mainly result from an averaging process of synaptic currents, both excitatory (glutamatergic) and inhibitory (GABAergic).

Therefore, LFPs may, in theory, give access to underlying synaptic currents and, subsequently, provide insights about the excitability level of recorded neuronal local networks. To reach this goal, the challenging issue is to estimate from a single extracellularly-recorded signal both ePSP and iPSP components of the LFP. In this paper, we propose a novel model-based approach to solve this problem. The method is based on model-based reverse engineering approach in which a neural mass model (NMM) is used to deconstruct the extracellularly-recorded field activity and reveal its main excitatory and inhibitory components. This reverse engineering method introduces two main steps. First, the classical NMM was revisited to solve an ill-posed problem, since two parameters (ePSPs and iPSPs amplitude) are estimated from a single signal (LFP recording) measured over a finite time window. Second, an optimization procedure was developed allowing for time tracking of ePSPs and iPSPs over a short-duration sliding window.

As demonstrated from both simulated and experimental data, the proposed approach can reliably estimate and monitor the average post-synaptic potentials and the densities of action potentials arising from underlying excitatory and inhibitory subsets of neurons. To our knowledge, this is the first study solving the inverse problem of LFP to estimate its main synaptic components based on a neural mass model. In a previous study by Einevoll et al. (Einevoll et al. 2007), the authors attempted to estimate synaptic activity from LFPs is already published using a laminar population analysis. Another related study by Gratiy et al. (Gratiy et al. 2011) estimated synaptic current source density with a model based on a single reconstructed cell morphology from the rat somatosensory cortex. One drawback of this method is that it requires anatomical information about cell morphologies or spatial distribution of synapses. However, those methods were applied on stimulus-evoked LFP responses in the rat somatosensory (barrel) cortex. Therefore, one motivation for our study was to develop a method that does not aim at reproducing stimulus-evoked LFPs, but rather spontaneous LFP signals from healthy/epileptic tissue, which can not be achieved by the aforementioned methods.

It is worth noting that several methods have been proposed to quantify the balance between excitation and inhibition (excitation to inhibition ratio, EIR). For instance, a relevant study used the slope of the power spectrum at low-frequencies (Gao et al. 2017) and identified that the latter was significantly associated with changes in the EIR. Despite its interest in terms of understanding how excitability shapes subtle features of the LFP power spectrum, this approach relies mainly of the power spectrum slope at low frequencies to estimate the EIR, and then excitatory/inhibitory components which are indirectly estimated through a multivariate regression model. Another study attempted to relate the EIR with the transfer function of a neural mass model, (Moran et al. 2007), but was an approximation using a linearized version of the system equation, without accounting for fine dynamics due to the hypothesis of stationarity. Our method overcomes these difficulties and provides a direct access to quantified indexes such as post-synaptic currents (excitatory/inhibitory, from which the EIR can be derived). Therefore, applications encompass diverse areas of neuroscience, ranging from basic neuroscience (tracking of excitability changes in neuronal networks) to clinics (analysis of depth-EEG recordings in patients with epilepsy).

## 2. Materials and methods

### 2.1 Principle of the model-based LFP reconstruction

The method exploits *a priori* knowledge about the processes (type and kinetics) underlying LFP generation in order to constrain LFP decomposition. LFPs are mostly composed of two sub-components (excitatory and inhibitory), and their extraction is challenging since 1) reconstructing two signals from only one is by definition an ill-posed problem; and 2) their frequency range is similar. In order to overcome this roadblock, we used a physiologically relevant NMM as a constraint to enable this decomposition process. NMM are a well-established computational class of models of neuronal population activity considering synaptic interactions between the neuronal assemblies involved in each LFP component: pyramidal cells (excitation) and interneurons (inhibition). We then used an LFP (experimental or simulated) to constrain the NMM response, and exploited this NMM to extract information contained in the LFP, especially to reconstruct excitatory and inhibitory processes.

### 2.2 Revised version of the neural mass model (NMM)

The NMM used in this study is an established and validated neural population model (Jansen & Rit 1995) including two sub-populations corresponding to pyramidal cells for excitation, and interneurons for inhibition. The corresponding block diagram of the model is presented in figure 1, where excitatory/inhibitory processes are represented in blue/green, respectively. In order to improve the physiological relevance of the NMM, we adapted the parameters of the “wave-to-pulse” sigmoid functions, converting the mean depolarization of sub-populations into an output firing rate, according to available neurophysiological data from the literature. We also reduced the number of parameters to be identified through the reverse modeling process by removing the connectivity parameters, which were instead integrated in the new sigmoid functions and in the synaptic gain parameters of the transfer functions. The main difference with the original Jansen-Rit’s model (Jansen & Rit 1995) is therefore the absence of connectivity parameters. Table 2 summarizes model parameters values. The resulting equations were derived following the method presented by Touboul et al. (Touboul et al. 2011). The synaptic impulse response functions used in the model were under the standard form: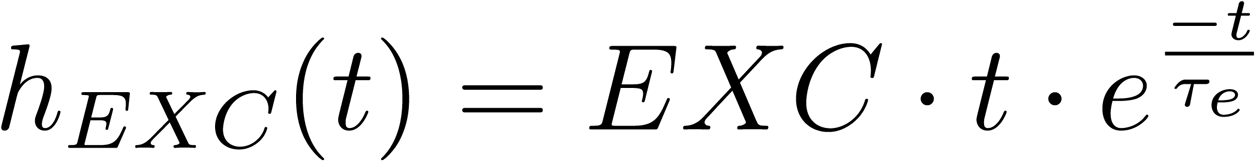 for excitation, and 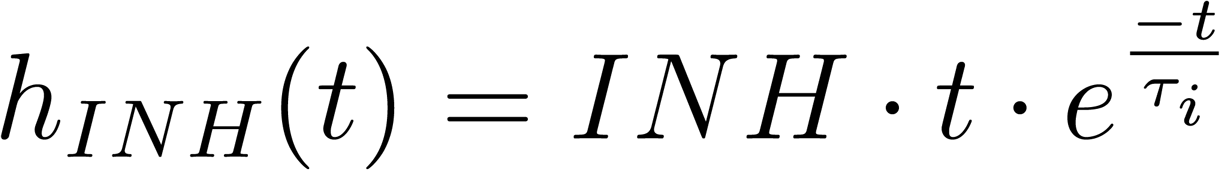 for inhibition, leading to a set of 6 differential equations forming the full NMM:

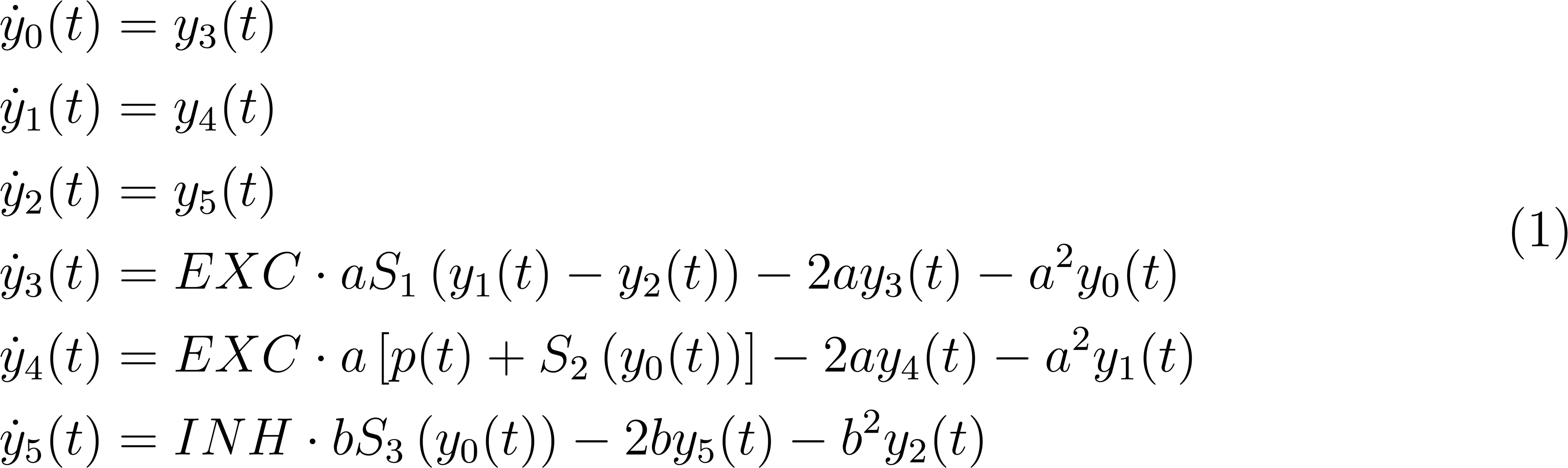

**Figure 1.**
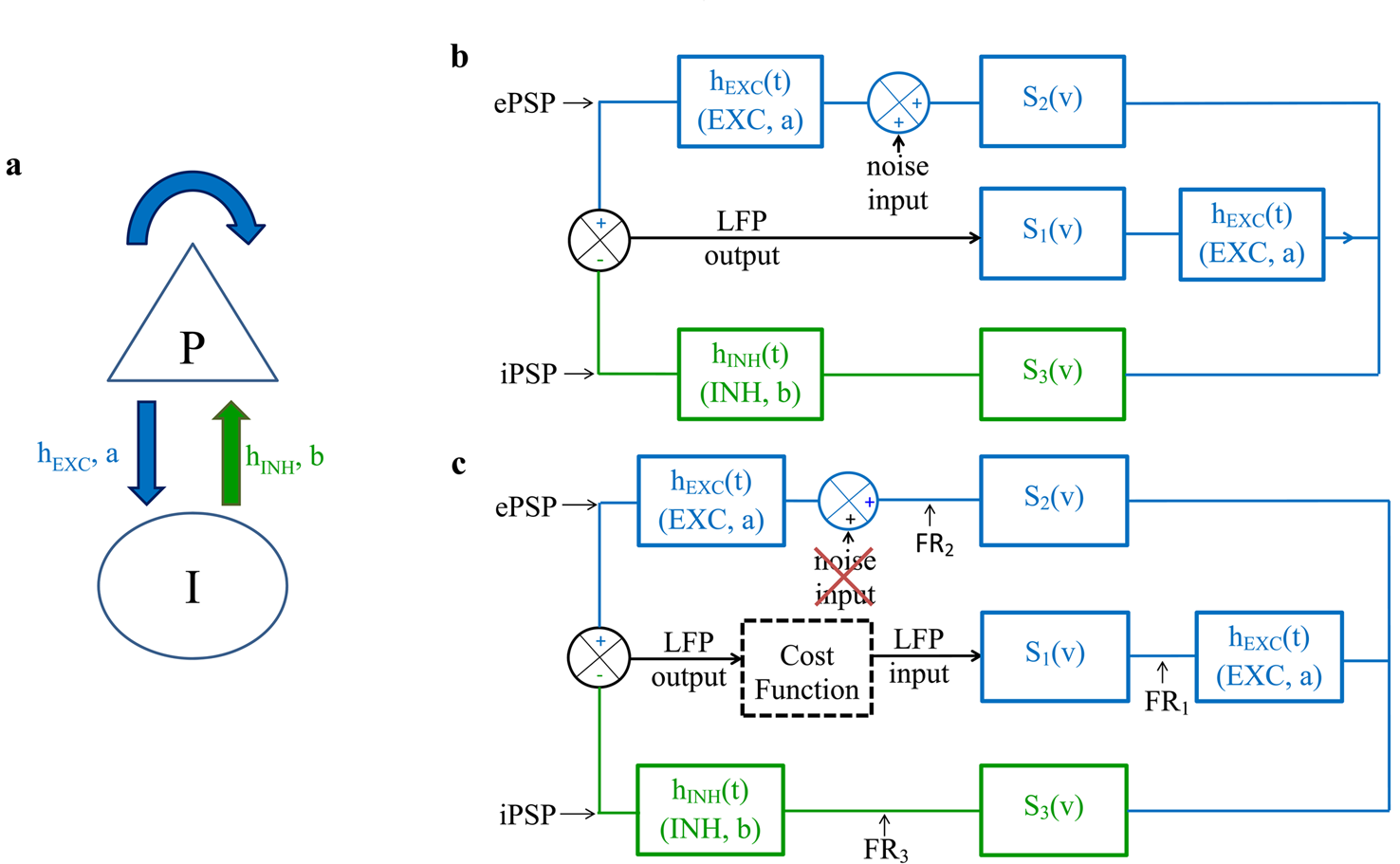
Illustration of the reverse modeling approach. (**a**) Neural mass model block diagram including two sub-populations: pyramidal cells (P) and interneurons (I). Excitatory (blue) and inhibitory (green) links between sub-populations are represented with arrows. (**b**) Neural mass model diagram where sigmoid functions were modified and connectivity parameters from the original NMM were removed. Excitatory and inhibitory loops are represented respectively in blue and green. LFP is the summation of synaptic inputs onto the pyramidal population (**c**) Reverse modeling approach where the new neural mass model is used to estimate PSPs contained in an LFP signal, the latter being set as an input of the model. A cost function based on the RMS error between estimated 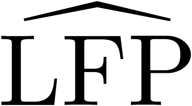 (LFP output) and input LFP was introduced to improve identification of EXC (excitatory) and INH (inhibitory) gain parameters. ePSPs: excitatory post-synaptic potentials, iPSPs: inhibitory post-synaptic potentials, *h*: synaptic impulse response functions, *S*_*n*_: sigmoid functions, 1*/a* and 1*/b*: dendritic average time constant.

Table 1 lists the studies that were used to identify sigmoid function parameters corresponding to pyramidal cells and interneurons. Since, in these studies, the pulse-to-wave relation was studied with respect to current, the first step was to convert the data with respect to voltage. To do so, a current-to-voltage conversion was used according to Destexhe and Paré’s study (Destexhe & Paré 1999), which used Ohm’s law with the membrane resistance. Then, a non-linear least square (Levenberg-Marquardt algorithm) was used to fit data from the literature, and an average was made between the two available sets to obtain sigmoid function parameters used in our model. The three sigmoid functions that were included in the NMM are:

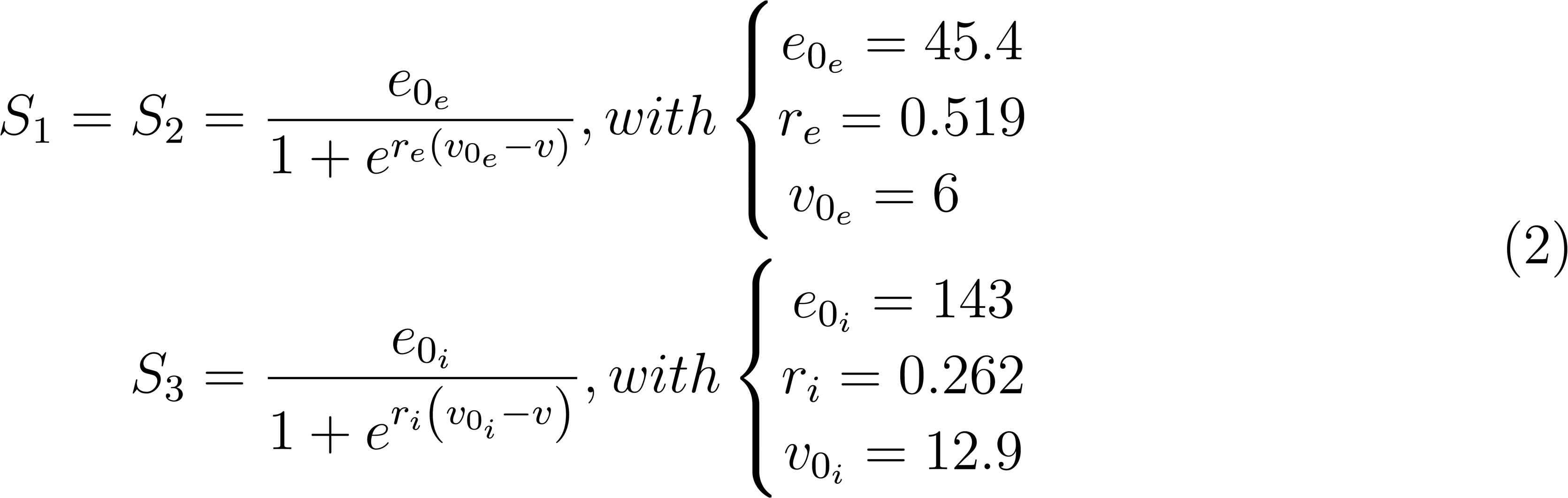

**Table 1.** Identified sigmoid function parameters.

| Pyramidal cells |  |  |  |
| --- | --- | --- | --- |
| References | Sigmoid function parameters |  |  |
|  | e0 | r | v0 |
| Reid et al. (Reid et al. 2014) | 48.9 | 0.534 | 5.12 |
| Chen et al. (Chen & Bazan 2005) | 26.7 | 0.721 | 5.81 |
| Malik et al. (Malik & Chattarji 2011) | 20.5 | 0.435 | 5.81 |
| Malik et al. (Malik & Chattarji 2011) | 112 | 0.356 | 6.62 |
| Wang et al. (Wang et al. 2015) | 19.1 | 0.550 | 6.67 |
| Mean \| | 45.4 | 0.519 | 6.00 |

**Table 1.** Identified sigmoid function parameters.
| Interneurons |  |  |  |
| --- | --- | --- | --- |
| References | Sigmoid function parameters |  |  |
|  | e0 | r | v0 |
| Gibson et al. (Gibson et al. 2006) | 173 | 0.166 | 17.7 |
| Ferguson et al. (Ferguson et al. 2013) | 128 | 0.239 | 12.0 |
| Fidzinsk et al. (Fidzinski et al. 2015) | 195 | 0.320 | 11.1 |
| Fidzinski et al. (Fidzinski et al. 2015) | 154 | 0.210 | 13.1 |
| Erisir et al. (Erisir et al. 1999) | 66.3 | 0.376 | 10.6 |
| Mean \| | 143 | 0.262 | 12.9 |

**Table 2.** Model parameters.

| Parameter | Interpretation | Value |
| --- | --- | --- |
| EXC | Average excitatory synaptic gain | 60 |
| INH | Average inhibitory synaptic gain | 15 |
| $\tau_e = 1/a$ | Dendritic average time constant<br>in the feedback excitatory loop | 1/100 s |
| $\tau_i = 1/b$ | Dendritic average time constant<br>in the feedback inhibitory loop | 1/35 s |
| $V_{0e}$<br>$e_{0e}$<br>$r_e$ | Parameters of the sigmoid<br>function for pyramidal cells | $V_{0e} = 6$ mV<br>$e_{0e} = 45.4$ Hz<br>$r_e = 0.519$ mV<sup>-1</sup> |
| $V_{0i}$<br>$e_{0i}$<br>$r_i$ | Parameters of the sigmoid<br>function for interneurons | $V_{0i} = 12.9$ mV<br>$e_{0i} = 143$ Hz<br>$r_i = 0.262$ mV<sup>-1</sup> |
| $p(t)$ | Excitatory input noise<br>(Gaussian white noise) | $m = 90$ Hz<br>$s = 30$ Hz |

### 2.3 Estimation of excitatory and inhibitory post-synaptic currents

We verified that this novel NMM could reproduce the four possible types of activity of the original model, namely: background activity, sporadic epileptic spikes, rhythmic spikes and narrow band activity at theta and alpha frequencies (see Figure S1 ActivityMap in Supplementary materials). The reverse modeling approach uses LFPs (simulated or recorded experimentally) that are injected in the NMM. The originality in our approach is to force the NMM state at the node of the associated block diagram (see figure 1(c)), corresponding to where LFPs are generated (”LFP input”), forcing other model components to use the LFP that has been injected as an input. By comparing the block diagrams in figure 1(b) and 1(c), the LFP input of the reverse modeling approach (figure 1(c)) corresponds to the LFP output of the NMM (figure 1(b)).

The injected LFP was first normalized to ensure that its magnitude was similar to simulated LFPs. The physical unit of LFPs, as well as ePSP and iPSP, is in terms of Volts, and the normalization allows not considering differences in hardware amplification gain use for recording LFPs. Furthermore, the normalization was done only once for the entire time course of the analyzed signal, even if the LFP was split into windows afterward as explained below. The reverse modeling approach involved removing the noise input (s=0) to avoid adding a stochastic component to the process, and also to correctly adapt 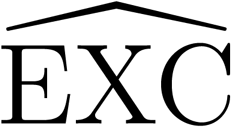 and 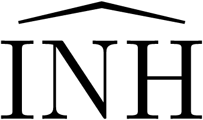 parameters to fit optimally the input LFP with the estimated 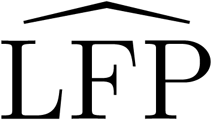. Since the model is then in an “open loop” form, the noise variance would just add noise onto ePSP. The method only allows the 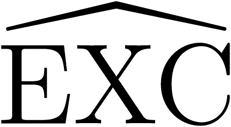 and 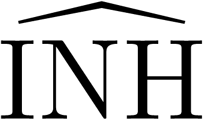 parameters to vary, since all other parameters relate to the intrinsic tissue properties. Therefore, the dendritic average time constants (1*/a* and 1*/b*) and the sigmoid function parameters (V_0e_, e_0e_, r_e_, V_0i_, e_0i_, and r_i_) were based on the literature. In order to ensure that NMM dynamics corresponds to the injected LFP, a key step consists in comparing the NMM output 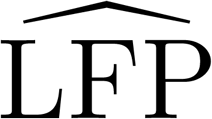 (estimated LFP computed in the reverse modeling approach) with the injected LFP, and to identify NMM gain parameters maximizing the match between LFP and 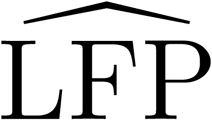. The identified gain parameters are 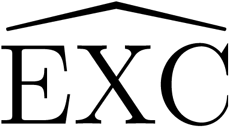 for the excitatory loop and 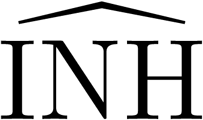 for the slow inhibitory loop. This identification was performed using a gradient descent, where the cost function was chosen as the root mean square (RMS) error between LFP and 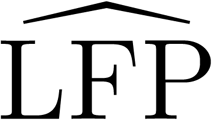 in magnitude (*M*_*RMS*_) and derivative (*d*_*RMS*_):

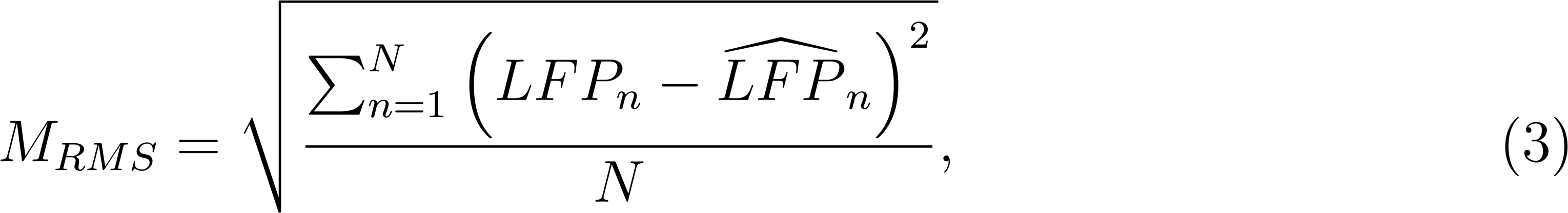

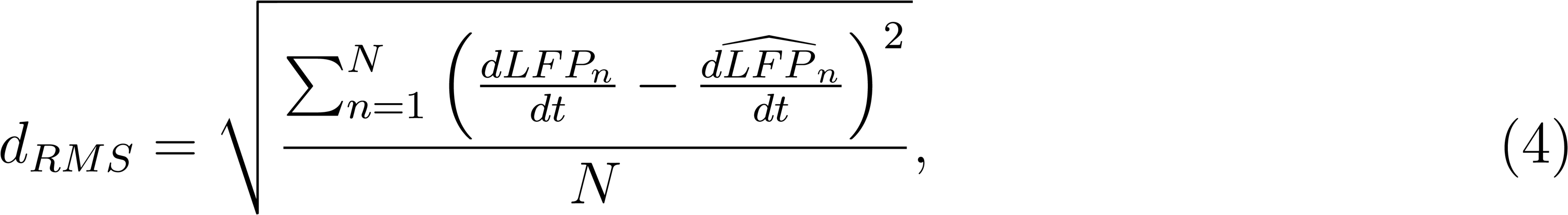

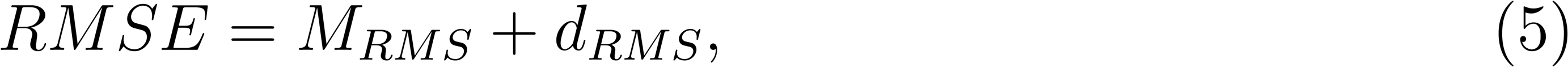

where *N* is the number of sample in the LFP.

Using both the difference in magnitude and derivative between LFP and 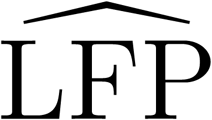 in the cost function enables evaluating the similarity in magnitude, but also in tendency. In the gradient descent method, both 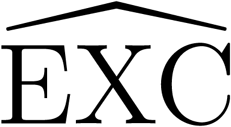 and 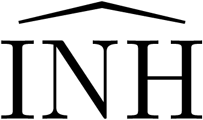 parameters were set to initial values (in a 2D parameter space) and the estimated 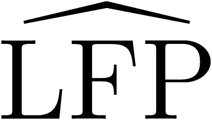 was computed for different set of parameters around the initial ones (on an ellipse where both radii were defined on the 2D parameter space *d*_1_ and *d*_2_). The gradient descent algorithm ended when no error lower than the current parameter set was found, and current 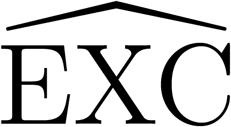 and 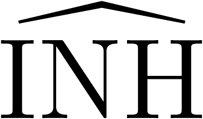 parameters were then taken as the optimal identification. It should be noted that this method does not identify all the basins of attraction in a 2D parameter space. As an attempt to verify that the set of identified parameters was unique, we also checked if there was, for a wide range of parameter values for EXC and INH, a unique solution under the form of a basin of attraction within the parameter space. This is a critical requirement for our method, since it would be problematic to have multiple potential parameter values for a single LFP to analyze. Parameters were set to form a 2D grid where EXC values varied from 0 to 100, and INH values varied from 0 to 50. The step for EXC was 0.25 and the step for INH was 0.125, leading to a 400×400 grid. For each parameter set, the RMS value between LFP and 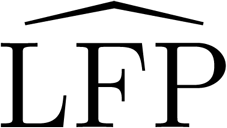 was computed. We present in figure 2 the corresponding result for qualitatively different LFPs (spikes, alpha, rhythmic and background activity), which highlight the uniqueness of the parameters set towards which the method converges. Note that this does not prove that it is impossible to have several basins of attraction, however we only identified one basin of attraction in all of our tests.

**Figure 2.**
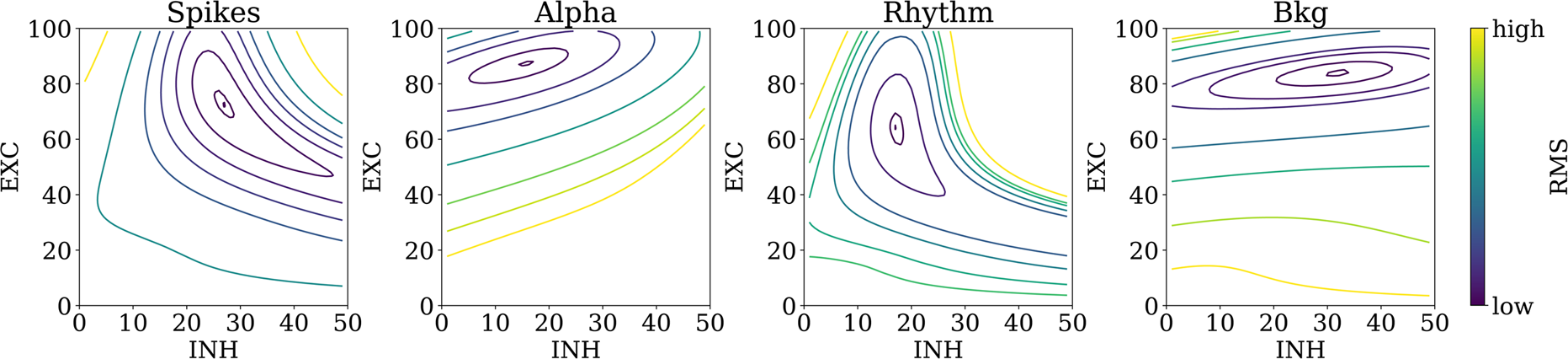
Illustration of the uniqueness of basins of attraction for parameter reconstruction using different LFP modes. RMSE computed within a 2D grid parameter space (EXC and INH). Note that only one basin of attraction is found for each type of activity.

Custom software was designed in Python, integrating a Qt interface (See Video S1 Software demo in Supplementary materials), dedicated to fitting at best NMM parameters on an injected LFP along short sliding windows on the time axis (i.e., NMM gain parameters are identified once per sliding window). Excitatory 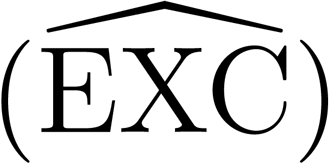 and inhibitory 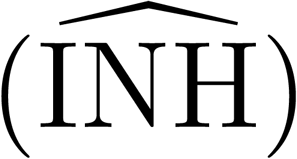 gains could therefore be tracked during the LFP time course, along with the excitation to inhibition ratio (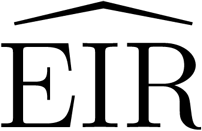, computed as the 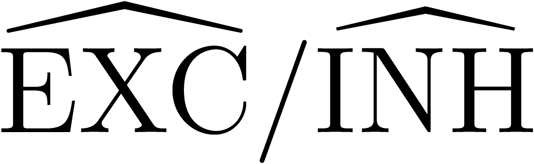 ratio). Obviously, the NMM provides full access at any time to the intermediate states for each sub-population, such as the average firing rate of action potentials and average post-synaptic potentials at the level of pyramidal cells and interneurons. Such signals are challenging to extract from LFPs with standard signal processing methods, especially since LFPs originate from the summation of PSPs onto pyramidal cells, and are not directly related with APs at the soma level. Using our approach, FRs can be directly deduced from a LFP with the reverse modeling approach, since FRs are intermediate states in the NMM.

### 2.4. Goodness-of-fit

The method used to compute the goodness of fit between an LFP and a reconstructed LFP was the zero normalized cross correlation (ZNCC (Nakhmani & Tannenbaum 2013)). This method provides an indication of similarity between two signals, named *γ*, which is bound [-1;1]. This indicator is 1 when both signals are identical, −1 when one signal is the opposite of the other, and is close to 0 when both signals are far from similar. This indicator does not depend neither on the length of the signals nor on the amplitude scales of different LFP measures (different amplifier gains for instance). Therefore, the ZNCC is especially suited to compare results for different LFP time/amplitude scales.

### 2.5. Simulated epileptic activity using another published model

In order to compare the performance of our model-based reconstruction of excitatory and inhibitory post-synaptic currents, we also used a previously published model simulating epileptic dynamics. The *Z*^6^ model (Koppert et al. 2016), based on the one in (Kalitzin et al. 2010), has only one parameter to control the excitability of the system (’c’). When −1 ≤ *c* < 0, this model generates steady-state behavior for values close to −1 (background activity), or a limit cycle for values close to 0 (ictal activity). All other parameters were fixed: a = −2; b = 2; *ω*_*mean*_ = 1; *ω*_*sd*_ = 0.2; *η*_*mean*_ = 0; *η*_*sd*_ = 0.2.

### 2.6. In vivo recordings

Electrophysiological recordings from N=7 mice were performed in accordance with the European Community Council Directive of November 24th 1986 (86/609/EEC), and were approved by the local ethics committee from the University of Rennes (agreement No 7872-2017031711448150). In all cases, bipolar electrodes (tips 400 *µ*m apart) were implanted bilaterally in the hippocampus (−2 mm anteroposterior, −1.5 mm mesio-lateral, −2 mm dorso-ventral from the Bregma) of C57B6j/Rj mice (80 days old). A reference electrode (monopolar) was placed at the cerebellum level. Mice were made epileptic following an intra-hippocampal injection of kainic acid that triggered an approx. 4-week epileptogenesis phase preceding the chronic epileptic phase. LFPs were sampled at 2048 Hz and hardware highpass filtered (0.16 Hz cutoff frequency). Recordings were made after the 4-week epileptogenesis period (chronic epilepsy stage). After protocol completion, mice were euthanized using the CO2 gradient method.

Electrophysiological recordings in rats (N=1) were obtained from the company Biotrial (http://www.biotrial.com). Ethics approval was obtained for these recordings, which involved injection of convulsive PTZ doses. A single bipolar electrode was implanted in the rat cortex, while a reference electrode was placed at the cerebellum level. A surgically implanted wireless transmitter enabled recording throughout the entire experiment without any physical manipulation of the rat, avoids movement artifacts. PTZ injection was performed through perfusion at the dose of 75 mg/kg.

### 2.7. In clinico recordings

In the context of pre-surgical planning, epileptic patients candidate to surgery undergo stereoencephalography (sEEG), involving the implantation of multiple intracranial electrodes aiming at improving the identification of epileptogenic zones. sEEG recordings from N=5 patients in the context of clinical evaluation were obtained from the Epileptology Unit of Rennes University Hospital (CHU Pontchaillou, Rennes, France). sEEG recordings were sampled at 2048 Hz with a Deltamed EEG acquisition system. Seizure epochs, involving several minutes pre- and post-seizures, were extracted for each patient to enable LFP reconstruction pre-, per- and post-seizure.

All the data are available on the following public link https://zenodo.org/deposit/1285630 and with the DOI: 10.5281/zenodo.1285630

## 3. Results

### 3.1 In silico validation

The reverse modeling method was first tested on simulated data. The NMM used in the reverse modeling algorithm used as an input the simulated LFPs generated by the same NMM. In this first validation step, a unique advantage is the ground-truth: estimated parameters (denoted by 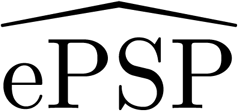 and 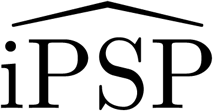) can be directly compared to actual components (ePSP and iPSP) of the simulated LFP used as an input signal.

Figure 3 presents the method performance on simulated data. Scatter plots in figure 3(a) display estimated parameter values (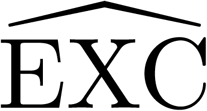, 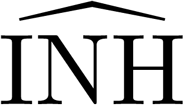 and 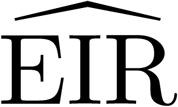: excitation to inhibition ratio) versus reference parameter values (denoted as EXC, INH and EIR) for a set of 35 simulated LFPs. The reverse modeling approach provides an excellent identification of parameters, since points in the scatter plot are close to the ideal result (represented by black lines when identified and reference parameters are equal). The method also provides an excellent 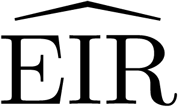 estimation, represented by the 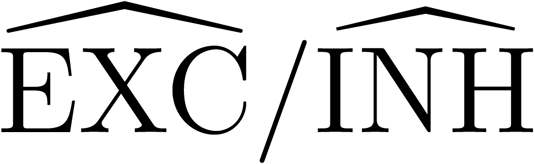 ratio. Panels in figure 3(b) to 3(d) present 4 qualitatively different types of LFPs (sporadic spiking, alpha, ictal and background activity) generated from the model, and the corresponding reconstruction computed from the proposed approach. Panels in figure 3(b) display LFPs, while panels in figure 3(c) display ePSPs, and panels in figure 3(d) display iPSPs. As depicted in figure 3(b), results revealed an excellent LFP reconstruction from estimated synaptic components, as shown by the goodness-of-fit indexes close to 1 (*γ* =0.999 for spikes, *γ*=0.962 for alpha, *γ*=0.993 for rhythmic and *γ*=0.909 for background). For background activity, the goodness-of-fit index was lower (*γ*=0.909) due to the strong contribution of the stochastic noise input (p(t)) for this particular type of activity. Reconstructed LFPs were obtained from estimated postsynaptic potentials, either excitatory 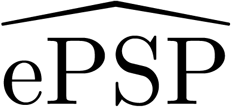 or inhibitory 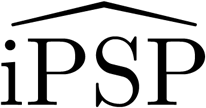, respectively shown in panels in figure 3(c) and 3d. As depicted, these estimated components perfectly fit the reference components (ePSPs and iPSPs) that were summed up to obtain the simulated LFP signal used as an input for the reverse modeling approach.

**Figure 3.**
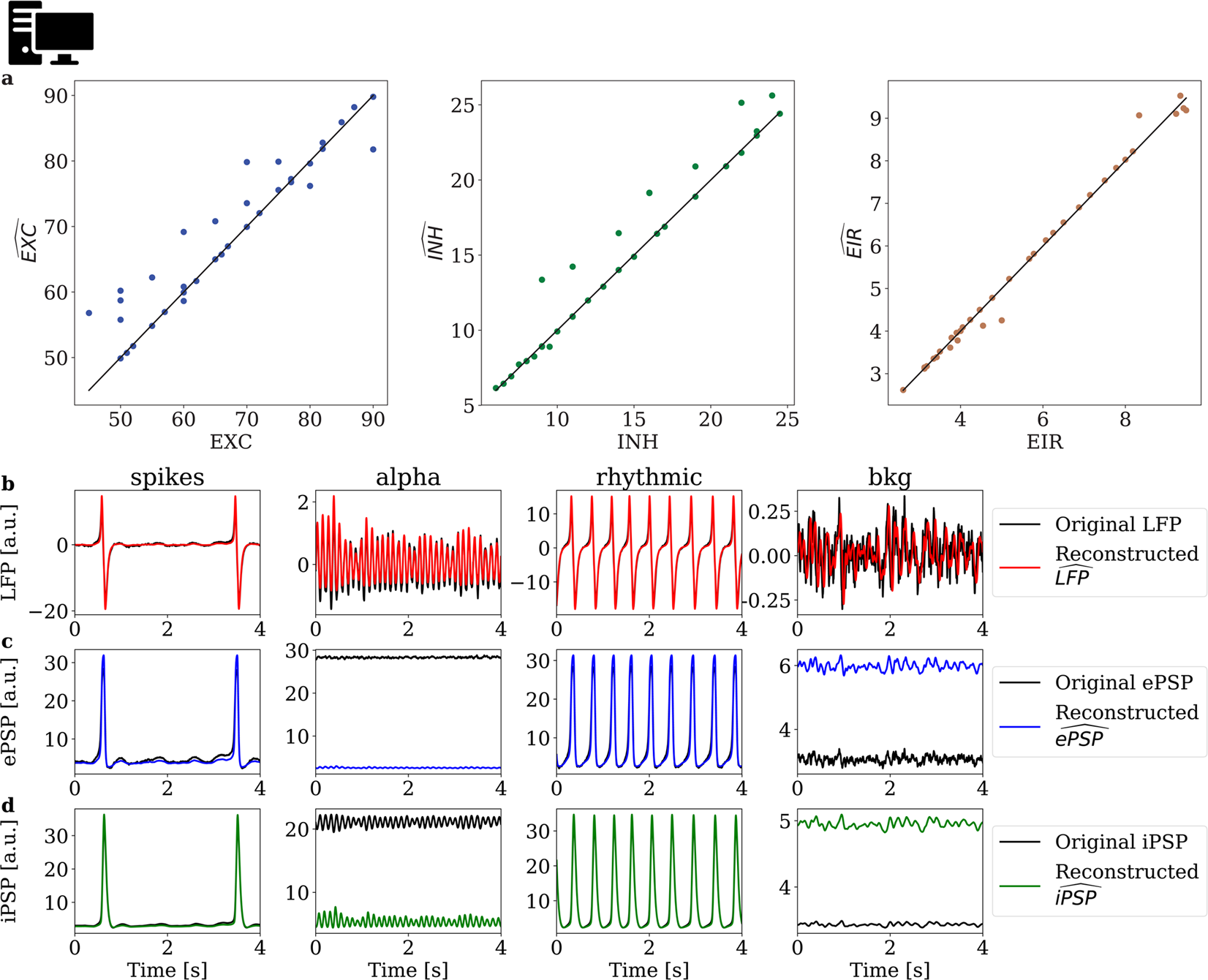
Results for simulated LFPs obtained using the same model as the reverse modeling approach. (**a**) Scatter plots for 35 simulated LFPs where ideal parameters vs identified parameters are displayed. Columns of combined panels b to e represent four types of activities. (**b**) LFP signals generated with the model (black) and 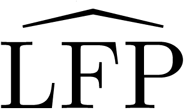 signals reconstructed with the reverse modeling approach (red). Goodness-of-fit indexes are: *γ*=0.999 for spikes, *γ*=0.962 for alpha, *γ*=0.993 for rhythmic and *γ*=0.909 for background (bkg), (**c**) ePSP signals generated with the model (black) and 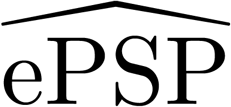signals extracted from the reverse modeling approach (blue). (**d**) iPSP signals generated with the model (black) and 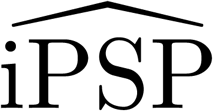 signals extracted from the reverse modeling approach (green).

In the previous example, the same model has been used both to generate LFPs and as the core for the reverse modeling approach. In order to perform a validation using another model based on a different formalism, we present below the results for LFPs generated with the *Z*^6^ model ((Kalitzin et al. 2010), see Methods section). We generated 150 s long LFP signals, with different values of the parameter *c* (from −1 to −0.01). Our method was applied on a 4 s windows with 1 s shift. The result is displayed in figure 4. The overall goodness-of-fit was *γ*=0.948, which shows an excellent agreement between the generated LFPs and reconstructed LFPs. Results point at a small decrease of 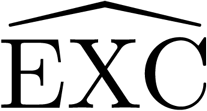 and a more pronounced decrease of 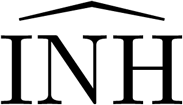 for increasing *c* values, leading to an EIR increase. Note that the 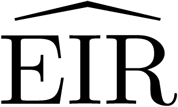 increases when the *c* parameter decreases, which in both cases results in an increase of excitability. In addition, our method also captures the non-linearity of the *Z*^6^ model, as shown in figure 4(c) where an exponential variation is seen. Since the *c* parameter controls the excitability of the system, we conclude that our method is able to capture the same excitability increase from LFP signals generated with another model of epileptiform activity. In other words, the performance of our method does not rely on the type of computational model used to generate epileptiform signals.

**Figure 4.**
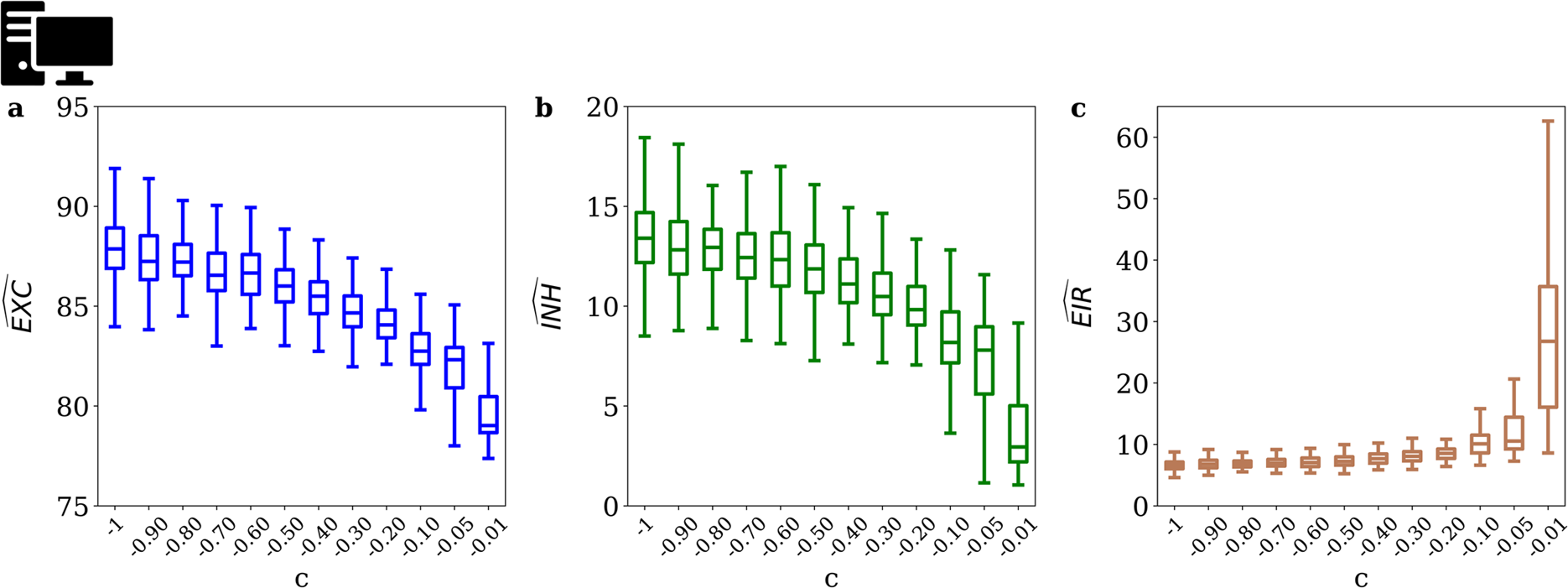
Results obtained with the *Z*^6^ model with different *c* values (duration: 150 s per LFP signals for each value of *c*). 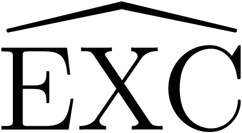 and 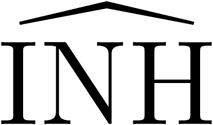 were identified using our reverse modeling approach. (**a**) Boxplot of 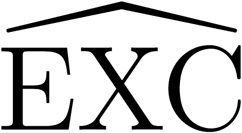, (**b**) 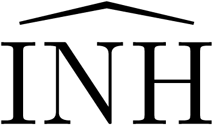, and (**c**) 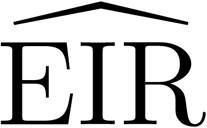.

In the following section, we move from the validation on simulated data to an experimental validation of the proposed method in the specific context of epileptic activity. This choice was motivated by the fact that the major qualitative changes observed in LFPs (during the interictal to ictal transition for example) involve drastic changes in neuronal excitability, potentially quantified by the proposed approach. Typically, seizures are associated with a significant increase of the excitation to inhibition ratio (Fritschy 2008).

### 3.2 In vivo and in clinico evaluation of the proposed approach

#### 3.2.1. PTZ-induced seizure (rat)

*In vivo* results obtained in rats following PTZ (pentylenetetrazol) intra-peritoneal injection are reported in figure 5. Results indicated that the proposed method leads to a reconstructed 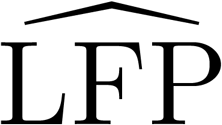 signal (red line, figure 5(a)) that was in excellent agreement (normalized goodness-of-fit between 0.85 and 0.97) with the experimentally-recorded LFP (black line, figure 5(a)) for the entire recording duration (130 seconds), as depicted on the 4 selected epochs (figure 5(a)). This illustrates the robustness of our estimation method, which provides accurate estimates for epochs featuring qualitatively different dynamics. In addition to close replication of LFP dynamics, the estimation of the excitation/inhibition ratio 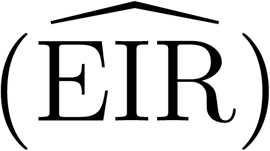 presented in figure 5(b), was in agreement with the expected excitability changes during the seizure time course: the 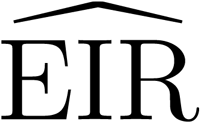 transiently increased at the onset of the PTZ-induced seizure, and then dramatically increased again during the seizure. It gradually decreased back to baseline as the seizure terminated. Interestingly, a decrease in the excitation level is observed during the seizure (figure 5(b)), which is however more modest than the drastic drop in inhibition, therefore still resulting in an increased EIR. Furthermore, since 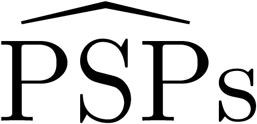 are computed in the NMM, they can be estimated through the propose method as presented in figure 5(c). The gradual decrease in inhibitory post-synaptic potentials associated with the seizure is clearly visible from epoch 1 to epoch 3, with a slight recovery in epoch 4 (figure 5(c)). Overall, there is satisfactory qualitative and quantitative agreement of LFP dynamics on the one hand, and of estimated 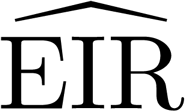 and 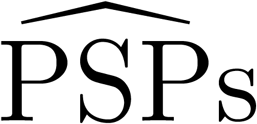 on the other hand. This supports the validity of our model-based reconstruction of excitatory and inhibitory components in data obtained *in vivo*. In addition, our method points at a result that is non-intuitive, namely the slight excitation decrease at the seizure onset.

**Figure 5.**
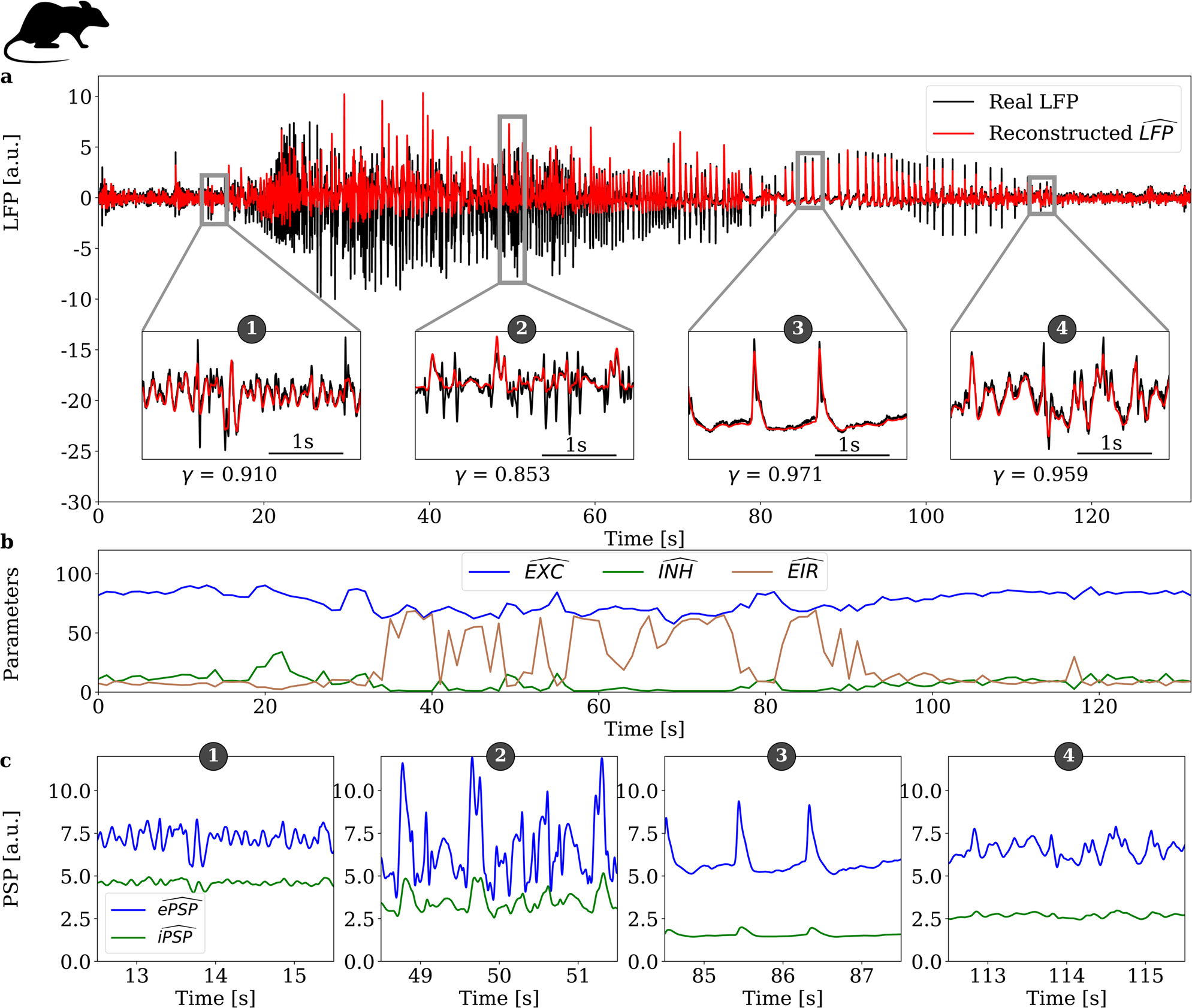
*In vivo* validation of the model-based LFP reconstruction in a PTZ-induced seizure (rat, N=1). (**a**) LFP signals recorded during a PTZ-induced seizure in rat (black line) and 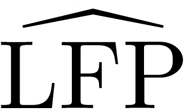 signals reconstructed using the reverse modeling approach (red line) for different types of activity (panels numbered from 1 to 4). The normalized goodness-of-fit index *γ* is presented for each activity type. (**b**) Identified model parameters: excitatory 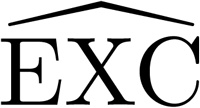 (blue line), inhibitory 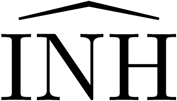 (green line), and the excitability ratio 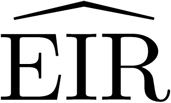 (brown line). (**c**) Reconstructed 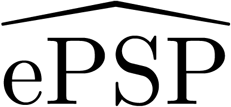 and 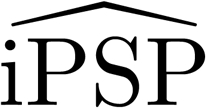 extracted from the NMM for each numbered panel presented in (**a**).

#### 3.2.2. Interictal activity in the kainate model of epilepsy (mouse)

The proposed method was also tested in epileptic mice (kainate model, see Materials and Methods section). Based on our method applied to 53 epochs of interictal events (hippocampal paroxysmal discharges, HPD) in N=7 mice, a reconstruction of excitatory and inhibitory components along with the 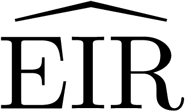 was performed. As depicted in figure 6(a), the model-based reconstruction of the 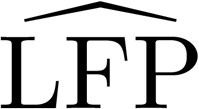 (red line) was in excellent agreement (goodness-of-fit index *γ* between =0.89 and 0.94) with the experimental LFP (black line) for the four considered epochs (see numbered panels in figure 6(a)). The reconstructed 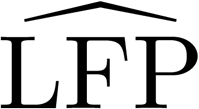 accurately matched the experimental LFP not only during background activity (panel 3 in figure 6(a)), but also during HPD (panel 4 in figure 6(a)). In addition, the time course of the estimated 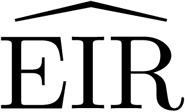 (figure 6(b)) was consistent with the occurrence of HPD within the signal: the 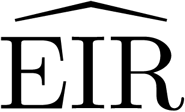 remained low during background activity, transiently increases during HPD, before returning rapidly to background values (figure 6(b)). Interestingly, as can be seen in figure 6(b), there was both a reduction of 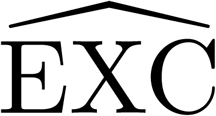 and 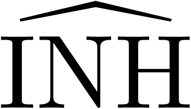 components (and not a decrease of 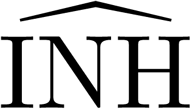 only as could be intuitively deducted in this case), which was however greater for 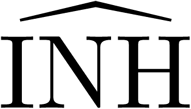, still resulting in a major 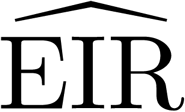 increase. This is similar to the results obtained in the case of the PTZ-induced seizure (see Section 3.2.1). In addition, figure 6(c) presents reconstructed post-synaptic potentials (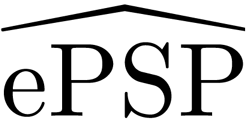 and 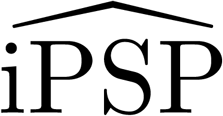) corresponding to each of the four LFP epochs in figure 6(a). The reconstructed iPSP points at a drastic inhibition decrease during a spike (epoch 2 in figure 6(c)) or HPD (epochs 1 and 4 in figure 6(c)), as compared to background (epoch 3 in figure 6(c)). The slight decrease in the reconstructed excitatory PSPs during interictal events (spikes or HPD) is also apparent (panels 1, 2 and 4 as compared to panel 3 in figure 6(c)). Figure 6(d) presents results at the group level (N=53 epochs) regarding the excitation and inhibition levels during HPD as compared to background activity epochs. We observed a significant decreased of excitation and inhibition (p *<* 0.001) during HPD as compared to background activity, with a lesser excitation decrease as compared to the inhibition decrease, which still results in a drastic 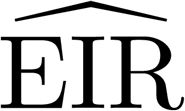 increase.

**Figure 6.**
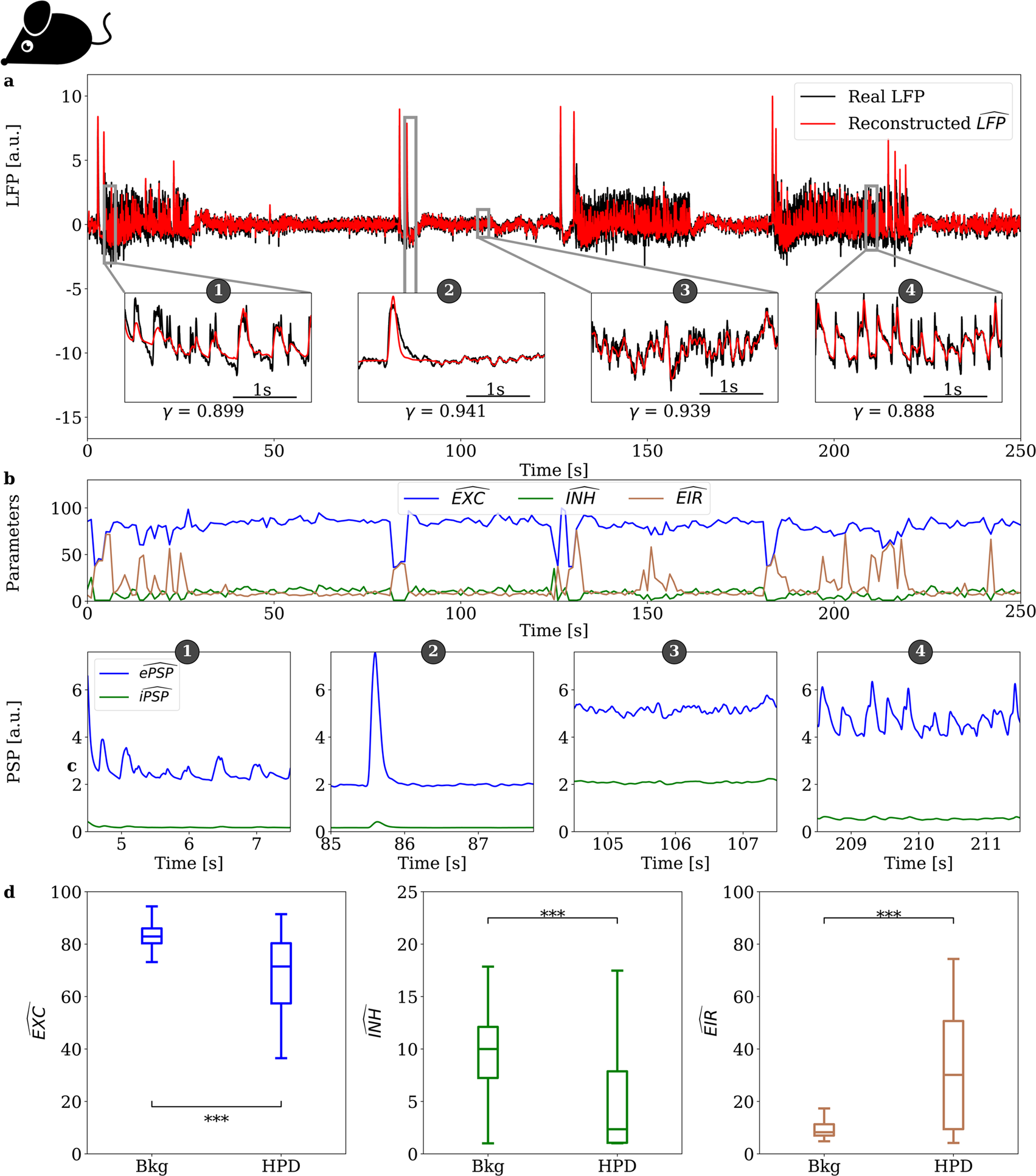
*In vivo* validation of the model-based LFP reconstruction in kainate mice (N=7). (**a**) LFP signals recorded in mouse where HPD are present (black line) and 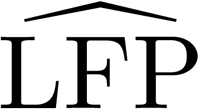 signals reconstructed with the reverse modeling approach (red line) for different types of activity (panels numbered from 1 to 4). The normalized goodness-of-fit index *γ* is presented for each activity type. (**b**) Identified model parameters: excitatory 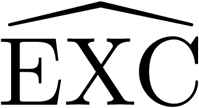 (blue line), inhibitory 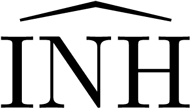 (green line), and the excitability ratio 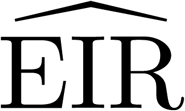 (brown line). (**c**) Reconstructed 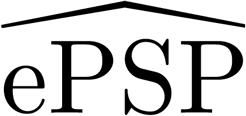 and 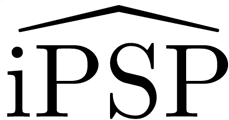 extracted from the NMM for each numbered panel presented in (a). (**d**) Boxplot of 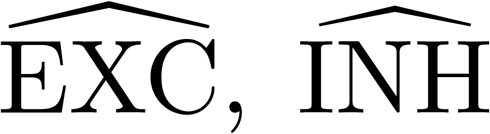, and 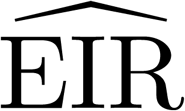 over background activity vs. HPD segments of 53 LFPs (*≈* 500s long) in seven mice. *\* * \** stands for p*<*0.001.

#### 3.2.3. LFP recordings after injection of a sub-convulsive dose of PTZ (mouse)

In the third *in vivo* experiment, a non-epileptic mouse was injected with a sub-convulsive dose of PTZ, aiming at validating our model-based method when LFP changes are subtler (i.e., where no seizures are present). The LFP recorded over the entire experiment (approx. 2200 seconds long) is presented in figure 7(a). Estimated 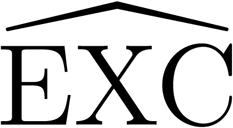 and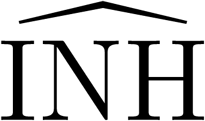 increase (p *<* 0.001) of the 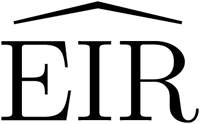 (figure 7(d)) shortly after injection, as compared to pre-injection levels. Consistently with the PTZ-induced rat seizure, the 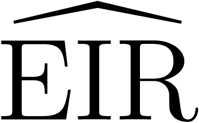 increased (figure 7(d)) due to a joint decrease of 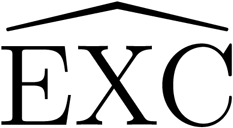 and 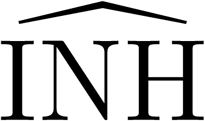 components, which is more pronounced for the 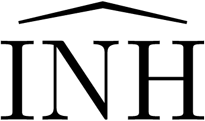 component than 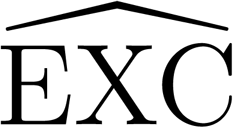. Furthermore, the 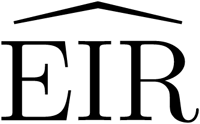 followed an exponential decay past the first 500 seconds after injection (the 500 s immediately after injection are discarded to avoid contamination by movements of the mouse due to manipulation by the experimenter), which is consistent with a pharmacokinetic response. The decay curve of the 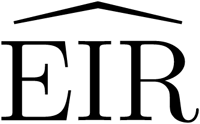 was fitted with an exponentially decaying curve, and the time constant optimizing the fit was estimated to be 370 seconds, in agreement with the PTZ pharmacokinetic time constant found in the literature (Wendling et al. 2016, Mandhane et al. 2007).

**Figure 7.**
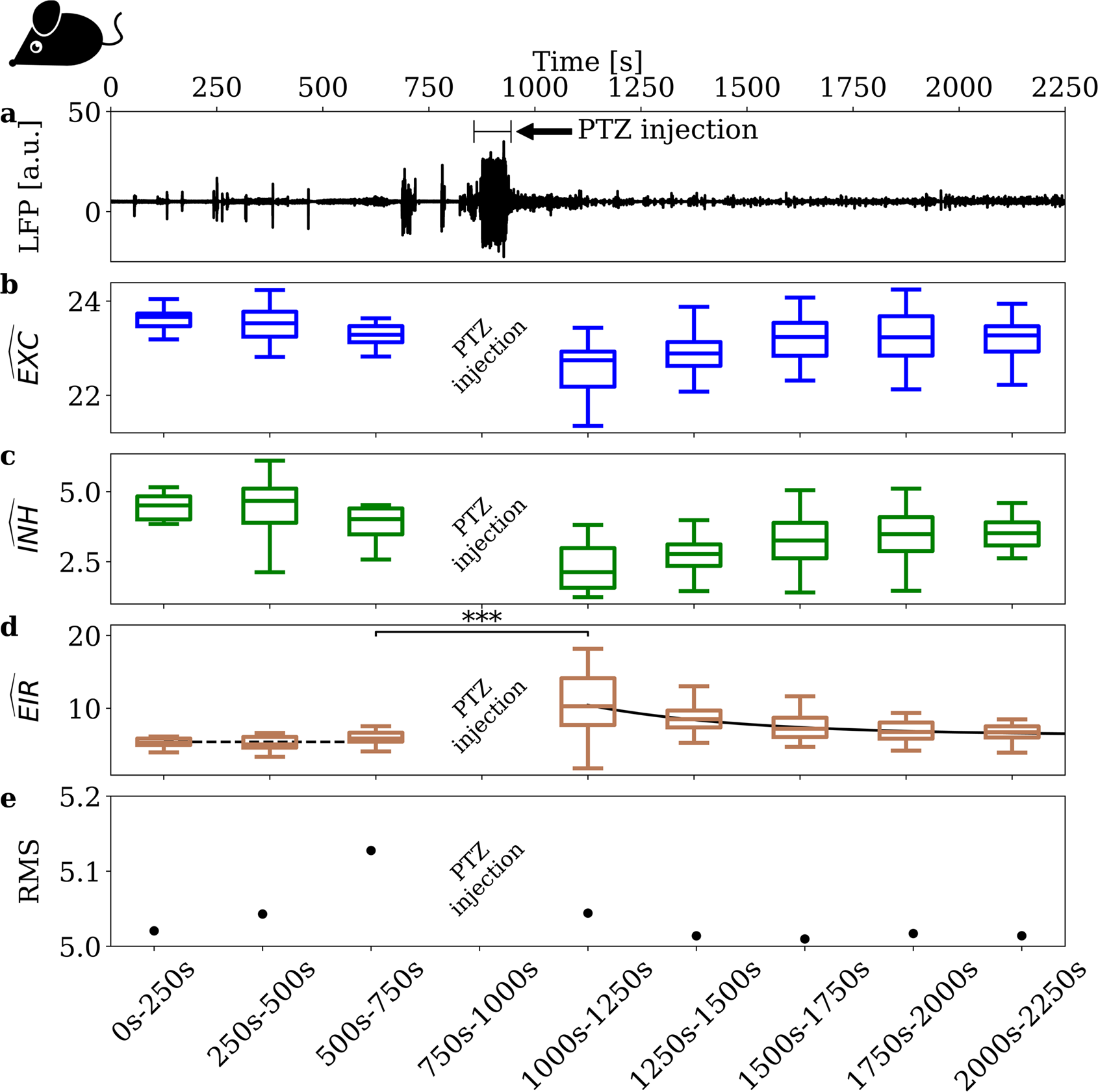
*In vivo* estimation of the excitation to inhibition ratio 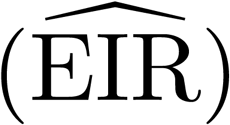 after injection of a subconvulsive dose of PTZ (mouse, N=1). (**a**) LFP recording. (**b**) Estimation of the 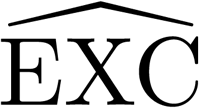 parameter. (**c**) Estimation of the 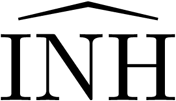 parameter. (**d**) 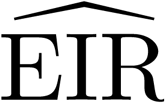. Dashed line corresponds to the average of three first boxplots. The continuous line corresponds to an exponential decay model (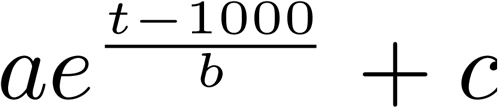, with a=4.1, b=-370 and c=6.3) on the five last boxplots. (**e**) RMS values computed for each time window. *\*\*\** stands for p*<*0.001.

#### 3.2.4. In clinico validation of the method in epileptic patients

Our method was validated using intracranial recordings from N=5 epileptic patients undergoing stereoencephalography (sEEG) in the context of pre-surgical evaluation. Among the available recordings, we extracted a total of 47 spontaneous seizures, which featured artifact-free epochs pre- and post-seizures, so that the 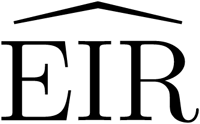 evolution could be tracked over the entire seizure time course. Figure 8 presents an example of human seizure and associated 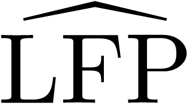 reconstruction (figure 8(a)) along with 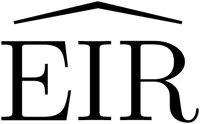 estimation (figure 8(b)), and reconstruction of PSPs (figure 8(c)). 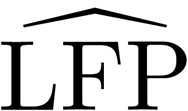 reconstruction is illustrated in figure 8(a), with the four zoomed panels emphasizing qualitatively different portions of the signal, illustrating the excellent agreement between the recorded intracranial signal (black line) and the reconstructed 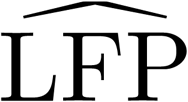 (red line), as quantified by the goodness-of-fit index ranging between 0.92 and 0.98. In the seizure presented as an example in figure 8(a), the estimated 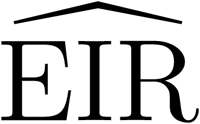 over the seizure time course (see figure 8(b)) was physiologically plausible, with a drastic 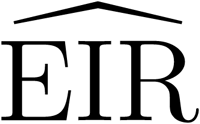 increase during the seizure itself, with a return to pre-ictal values after seizure cessation. The reconstructed post-synaptic components (figure 8(c)) for each of the four panels presented in figure 8(a) also pointed at a gradual decrease in the inhibition level as the seizure progresses, consistently with the *in vivo* PTZ-induced seizure studied (figure 5). Figure 8(d) presents the group results (N=5) and compares reconstructed excitation and inhibition levels during interictal and ictal epochs. The estimated 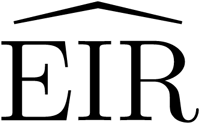 increase during seizure was driven by a significant 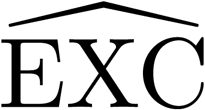 increase (p *<* 0.001), combined with a significant 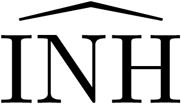 decrease (p *<* 0.001). Reconstructed 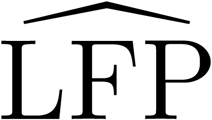s for this database matched closely experimental LFPs, with an average goodness-of-fit *γ*=0.893 *±*0.56.

**Figure 8.**
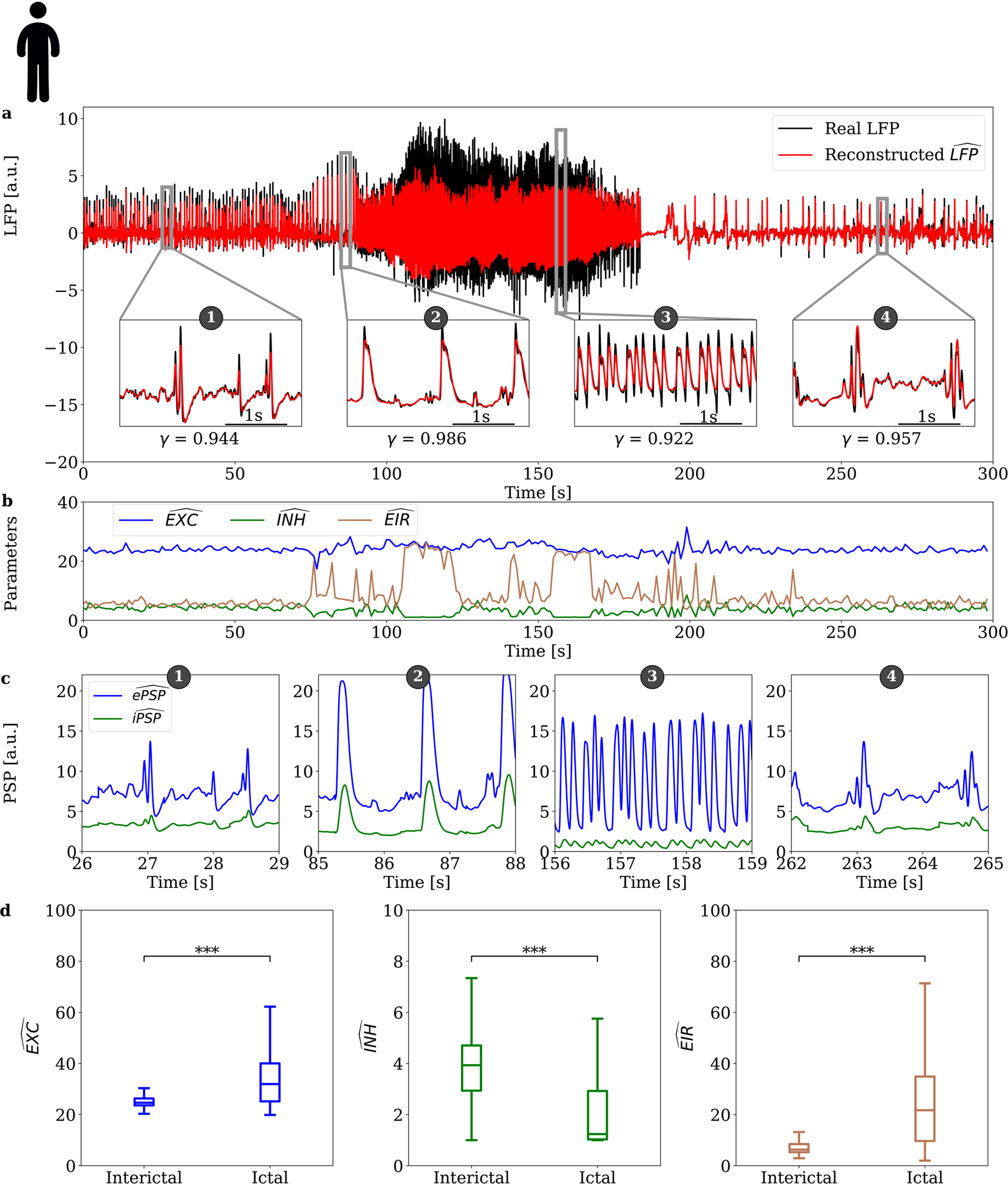
Validation on human seizures data (epileptic patients, N=5) obtained through intracranial recordings. (**a**) LFP signals recorded in human where a seizure is present (black line) and 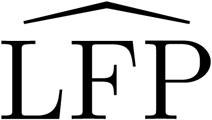 signals reconstructed with the reverse modeling approach (red line) for different types of activity (panels numbered from 1 to 4). The normalized goodness-of-fit index *γ* is presented for each activity type. (**b**) Identified model parameters: excitatory 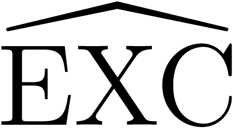 (blue line), inhibitory 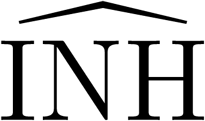 (green line), and the excitability ratio 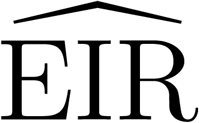 (brown line). (**c**) Reconstructed 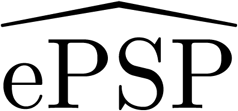 and 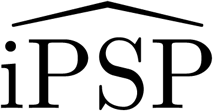 extracted from the NMM for each numbered panel presented in (a). (**d**) Boxplot of 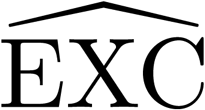, 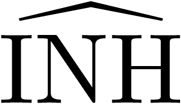 and 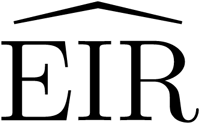 ratio over interictal vs ictal segments of 47 LFPs in five human patients. * * * stands for p<0.001.

## 4. Discussion

We developed a novel model-based method enabling the plausible reconstruction of excitatory and inhibitory post-synaptic currents involved in LFP generation as measured experimentally by intracranial electrodes. The method is based on a neural mass model, since this modeling approach captures crucial physiological components such as the dendritic average time constants, or wave-to-pulse sigmoid non-linear functions. While obviously such models do not capture all physiological aspects (e.g., neuron orientation with respect to the recording electrode as mentioned in (Einevoll et al. 2013) or cell morphology (Gratiy et al. 2011)), it stills features the two main excitatory (Glutamate) and inhibitory (GABA) post-synaptic currents, which is generic and widespread among cortical structures. Furthermore, this model is able to generate LFP signals with only a limited number of parameters. This method was tested at several levels: from *in silico* (using the method on LFPs generated using a NMM, providing a ground truth, as well as LFPs generated with another epileptic model, the *Z*^6^ model), to *in vivo* (rat and mouse LFPs), and finally to *in clinico* (sEEG recordings in epileptic patients). In both *in silico* tests, the method was able to track excitability according to EXC and INH for NMM-based LFPs, and the *c* parameter for *Z*^6^ model based LFP. In all model-based LFP reconstructions based on experimental data, we found plausible and relevant results emphasizing an 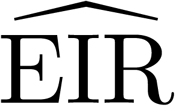 increase during the generation of interictal events and during seizures. Furthermore, the reconstructed 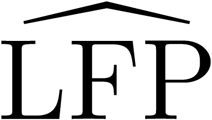, using our model-based decomposition, matched accurately experimentally recorded LFPs, suggesting that the assumption that the LFP is mostly generated by excitatory and slow inhibitory post-synaptic currents is reasonable. In addition, using a subconvulsive dose of PTZ in mice, we were able to provide an estimate of the PTZ time constant consistent with the literature. Therefore, a major advantage of our technique is that it enables the detection of subtle variations in neuronal excitability; such as those induce by sub-convulsive doses of PTZ. In addition, it is worth noting that the proposed method consistently pointed at a counter-intuitive decrease of excitation (minor as compared to the decrease of inhibition) in all *in vivo* recordings (PTZ-induced seizure, epileptiform events in the kainate model). However, we did not identify a similar decrease of excitation during human seizures from our available database. A possible explanation is that slightly different pathophysiological mechanisms are involved in the generation of epileptiform activity in our *in vivo* models as compared to human seizures.

The possibility to reconstruct, from an experimentally recorded LFP, the time course of excitatory (glutamatergic) and inhibitory (slow GABAergic) post-synaptic contributions, is especially appealing. Our method provides indeed full access to fundamental variables such as ePSPs and iPSPs for each subpopulation. These signals are challenging to extract from LFPs, since the problem is underdetermined. The identification of NMM parameters to optimize the fit of simulated LFPs with experimental LFPs has been addressed in several studies, using methods based on genetic algorithms (Wendling et al. 2005), Kalman filter (Bellanger et al. 2005), synaptic conductance estimation (Rudolph et al. 2004), or Hodgkin-Huxley parameters estimation (Zheng et al. 2012). However, none of these methods provides access to PSP signals from experimentally recorded LFPs. Our method should have applications in numerous areas of neuroscience, since LFP is a very common electrophysiological recording modality, from *in vivo* to *in clinico*, and ranging from neurophysiology to diagnostic applications. In addition, the method is sensitive enough to capture subtle changes in excitability, as shown in the case of the sub-convulsive PTZ dose experiment, and was able to estimate successfully PTZ pharmacokinetics.

One limitation is that our method currently features only slow, dendrite-targeting GABAergic synapses, and neglects fast soma-targeting synapses; which limits the possible frequency range of LFPs that can be reconstructed. This simplification was done for three reasons. First, we aimed at providing a proof-of-concept that a method based on a physiologically-based NMM can provide meaningful excitatory and inhibitory components from an experimentally recorded LFP. Second, due to its very generic structure, the NMM used should be general enough to analyze LFPs from any brain region. Third, if a sub-population of fast GABAergic neurons was added in the model, that would considerably increase the time required for parameter identification, since that would involve a 3D parameter space instead of 2D. This absence of fast GABAergic activity explains why the epochs where higher frequencies (beta band −13 to 30 Hz- and above) were present in the LFP are more challenging to reconstruct using our approach. This specific point explains why the goodness-of-fit is better in the case of HPD in mice (figure 6) as compared to epochs of relatively fast activity during the rat PTZ-induced seizure (figure 5): HPD are mainly characterized by slow activity, as compared to the relatively fast activity occurring at the onset of seizures. Therefore, agreement between reconstructed and experimentally measured LFPs is lower when higher frequencies (beta range and above, i.e. *>* 13 Hz) are present in the signal.

Using our model-guided approach to unveil the time course of excitatory and inhibitory post-synaptic currents is promising, especially since the algorithm provides excitability tracking with a performance in real-time (Video S1 Software demo in Supplementary materials): on average, 2 s of LFP sampled at 1024 Hz requires 0.7 s using a single core on a PC equipped with an Intel Xeon E5-2637 3.5 GHz CPU and 64 GB of 1866 MHz RAM. Furthermore, there is still ample room for computation time optimization, since the algorithm is currently programmed in Python, which is known for not being optimal in terms of computation time. The possibility to automatically track neuronal excitability in real-time could be implemented on-chip, for example, in a neuromodulation device, to trigger stimulation when excitability exceeds a certain threshold. Another immediate application would be the monitoring of epileptic patients undergoing sEEG for pre-surgical evaluation.

## Author Contributions

Conceived and designed the experiments: P.B., F.W. Computational model: M.Y., J.M. F.W. Performed the experimental study: P.B., J.M. Manuscript preparation: M.Y., P.B., J.M., F.W.

## Acknowledgment

This work is supported by NIH. Application Number: R01 NS092760-01A1.

